# Membrane dynamics of Notch-bound γ-secretase produces two distinct Notch conformations

**DOI:** 10.1101/607846

**Authors:** N. Tang, B. Dehury, K. P. Kepp

## Abstract

Cleavage of Notch by the major intramembrane aspartyl protease complex γ-secretase is a central event in cell regulation and is also important to Alzheimer’s disease, with more than 200 mutations in the catalytic subunit of γ-secretase (PS1) causing severe early-onset forms of the disease. Recently, cryogenic electron microscopy (cryo-EM) has revealed the electron density of the protein-Notch complex in frozen solution, indicating major changes upon substrate binding and a possible helix unwinding to expose peptide bonds. In order understand the all-atom dynamics that cause this process, and to test the Notch binding in a membrane protein rather than solution, we developed an all-atom model of mature wild-type γ-secretase bound to Notch in a complete membrane-water system and studied the system using three independent 500-nanosecond molecular dynamics simulations. Our ensembles are in essential agreement with known cryo-EM data. As in previous simulations we find unusual β-strand transitions in exposed parts of PS1. We also observe the atomic helix motions that cause loss of helicity in bound Notch by direct comparison to corresponding 500 ns simulations of free Notch, in particular five residues to the N-terminal site of the primary cleavage site. Most importantly, we identify three conformation states, with two of them differing in the Notch-bound catalytic site. These dynamics produce a ping-pong relationship of positioning the S3 cleavage sites of Notch relative to the aspartates. These conformation states are not visible in the cryo-EM data; probably the density is an average snapshot of the two states. Our identified conformation states rationalize how Notch cleavage can be imprecise and yield multiple products. Our identified conformation states may aid efforts to develop conformation-selective drugs that target C99 and Notch cleavage differently.

**Statement of Significance:** The atomic dynamics underlying cleavage of Notch by γ-secretase in the membrane is of major biological importance. Electron microscopy has revealed the protein-Notch complex in frozen solution, showing major changes upon substrate binding and helix unwinding to expose peptide bonds, but does not explain why substrate cleavage is imprecise and produces several products. Our model of wild-type γ-secretase bound to Notch in a complete membrane-water system equilibrated by 3 × 500 nanoseconds of molecular dynamics strongly complements the electron microscopy data: We identify the specific loop and helix motions that cause the β-strand transitions in PS1 and the loss of helicity in specific residues of bound Notch. We identify different conformations of Notch, which importantly affect the S3 cleavage site; the open state may cause the imprecise cleavage with earlier release of products. Our identified states can aid development of conformation-selective drugs that target C99 and Notch cleavage differently.

## Introduction

Alzheimer's disease (AD) is a major neurodegenerative disease causing gradual loss of memory, cognition, identity, life quality, and autonomy, ultimately leading to death, and affecting tens of millions of people worldwide. Despite its prevalence and devastating outcome, the complexity of the disease has prevented the development of causal treatments, and currently marked drugs only delay disease progress by months.(1–3) Recent failures in clinical trials of major drug candidates have led to calls for alternative and more detailed causative disease mechanisms.(4–8)

Senile plaques in patient brains consisting of aggregated β-amyloid (Aβ) is a central pathological hallmark of AD.(9–11) The Aβ peptides are formed by sequential cleavage of β-amyloid precursor protein (APP), first by β-secretase which removes the ectodomain and produces C99, and then by the catalytic presenilin (PS) subunit of γ-secretase, which trims C99 to Aβ within the membrane.(12–14) Due to the consecutive trimming of C99, the released Aβ peptides differ in size.(12, 15, 16) The shorter Aβ peptides, notably Aβ_38_ and Aβ_40_, are thought to be benign, whereas longer isoforms such as Aβ_42_ and Aβ_43_ are more hydrophobic, aggregation-prone, membrane-interacting, and more toxic to cells.(11, 17, 18) Genetic mutations in the genes coding for APP, PS1, and PS2 cause particularly severe early-onset familial AD, which supports the involvement of Aβ in the disease even beyond its presence in plaques.(19–22) The most persistent feature of these mutations is an increase in the Aβ_42_/Aβ_40_ ratio resulting from C99 cleavage,(12) and this ratio correlates directly with the clinical severity of mutation implying its clear relationship to disease.(23–26)

For these reasons, γ-secretase is currently explored as a major drug target.(27, 28) γ-Secretase is a large intramembrane aspartyl protease consisting of four subunits: PS1 or PS2, presenilin enhancer 2 (PEN-2), anterior pharynx defective 1A or 1B (APH-1A, APH-1B), and nicastrin (NCT). The catalytic subunit PS1/PS2 is comprised of nine transmembrane helices (TM1-TM9) and harbors the two catalytic aspartate residues (Asp257 in TM6 and Asp385 in TM7) which constitute the active site of γ-secretase.(14, 29, 30) Assuming that inhibition of γ-secretase activity would lower Aβ production and thus be beneficial for AD, a large number of γ-secretase inhibitors have been developed.(27, 31, 32) Unfortunately, none of them have shown significant endpoints, and side effects were observed in the clinical trials.(4)

The failure of the γ-secretase inhibitors can be due to several reasons: One is that Aβ should not be reduced indiscriminately because it serves important functions in the brain;(33) another is that inhibition of γ-secretase prevents the cleavage of perhaps hundred other substrates of this major membrane protease, including the systemically important Notch.(34–37) Notch is involved in multiple cell differentiation processes and plays an important role in neuronal function and development.(38, 39) Cleavage of Notch by γ-secretase occurs in a manner similar to C99(40) and releases the Notch intracellular domain (NICD), which plays an important role in learning and memory.(39, 41) As is the case for C99, Notch-cleavage can occur at multiple positions around the S3 site near the intracellular membrane border, but most of the resulting fragments are rapidly degraded according to the N-rule.(41) Non-selective γ-secretase inhibition will abolish the NICD generation and can contribute to cognitive decline.(42) Realizing the problems of γ-secretase inhibitors, Notch-sparing γ-secretase modulators have become a research focus in recent years.(5, 43) However, the potential molecular mechanism of such γ-secretase modulators is still not understood, and this limits their further development.(44)

The recently solved cryo-electron microscopy (cryo-EM) structures of γ-secretase bound to the shorter C83 analog of C99 and Notch-100 reveal the probable mode of Notch binding by γ-secretase, but also indicate how it differs from the corresponding binding of C99.(45, 46) These structures were solved without the membrane at cryo temperature, and the detailed atomic dynamics that produce the average density maps are thus not available. Most importantly, while insightful in many ways, the average structure does not explain why Notch (and C99) cleavage is imprecise or “sloppy”, leading to multiple cleavage products at the S3 site.(41)

To understand the underlying atomic dynamics of the Notch-bound γ-secretase in a realistic membrane at ambient temperature, we performed molecular dynamics (MD) simulations of an essentially complete (except the elusive N-terminal of PS1) all-atom model of mature auto-cleaved (as required for activity) wild type γ-secretase and the Notch transmembrane domain (TMD) in a membrane-water system. The computations were performed in 2018 before the cryo-EM structures became available and its general agreement with the cryo-EM structure thus testifies to the accuracy of our MD protocol, but provide the atomic motions that rationalize the cryo-EM data, and also show important differences that are not seen in the cryo-EM data.

Most importantly, the cryo-EM data do not allow the identification of several substrate-bound conformation states. Our membrane dynamics identify three states, with two of them distinctly affecting the substrate binding pocket, creating an open or a compact state upon Notch binding. These identified states explain why substrate processing by γ-secretase can be imprecise and lead to different-length NICDs.(41, 47)

## Computational methods

### Protein-protein docking

The previously published NMR ensemble of the Notch TMD (PDB: 5KZO)(48) was used to construct Notch models in the present study. Ten structural models were relaxed using Rosetta (version 2017.36), generating 20 minimized structures for each NMR model. The structure with the lowest energy was then selected for docking into γ-secretase. As starting model for γ-secretase, we used our previously established mature apo-structure.(49) In this structure, the hydrophilic loop 2 (HL2) has undergone autocleavage to obtain the mature, active state of the protein which has two PS1 fragments, the C-terminal and N-terminal fragments (NTF and CTF). Understanding how this maturation is required for activity(50) by changing the loop motions is essential to understand γ-secretase activity but these loop parts are not visible in the cryo-EM data and thus requires MD simulation as done here. This starting choice is partly due to our work being carried out during 2018 before the new structure was available, but importantly this choice also prevents bias toward the new substrate-bound cryo-EM structures and enables a strict test of our model-building protocol. The protein-substrate docking was performed using the PIPER method of the ClusPro webserver.(51) The top ranked structure with most clusters was used to build our model.

### Molecular dynamics simulations

The obtained model of the γ-secretase-Notch complex was embedded in a complete all-atom membrane model using the position protein in membrane (PPM) server with the original coordinates transformed to the middle of membrane.(52) The resulting structure with membrane boundary planes was then uploaded to the CHARMM-GUI web server to build the bilayer system.(53, 54) A homogeneous 1-palmitoyl-2-oleoyl-glycero-3-phosphocholine (POPC) lipid bilayer with a total of 350 lipids (175 per leaflet) was used, and the water thickness above or below the γ-secretase-Notch complex was maintained at 2 nm. In order to simulate a physiologically relevant system, because ion strength can affect protein conformations,(55) 173 Na^+^ and 158 Cl^−^ ions were randomly placed to obtain a realistic ionic strength of 0.15 M, using Monte Carlo randomization.

Using the resulting protein-Notch-membrane system as starting structure, MD simulations were performed with the GROMACS package (2018.2 GPU version).(56) The structure-balanced Charmm36m force field with compatible membrane-lipid parameters was used for the simulation.(57) The water was described using the TIP3P explicit 3-site water model.(58) Non-bonded interactions were evaluated until a cutoff of 1.2 nm using a Verlet cutoff scheme. The particle mesh Ewald (PME) algorithm was used for calculating long range electrostatic interactions.(59) The linear constraint solver (LINCS) algorithm was used to constrain the bonds connected to hydrogen atoms.(60)

The established system was first minimized using the steepest descent algorithm, followed by a six-step position restrained equilibration for 500 ps with the first two steps equilibrated in a canonical (NVT) ensemble and the next four steps equilibrated in an isothermal isobaric (NPT) ensemble using Berendsen temperature and pressure coupling, respectively.(61, 62) After this equilibration, three independent 500-nanosecond (ns) production simulations with a 2 fs time step were performed in an NPT ensemble using the Nose-Hoover thermostat and Parrinello-Rahman barostat at 303.15 K and 1 atm.(63, 64) The simulations were started using different velocity seeds made from a random number generator, to make them statistically independent. Based on previous studies of the apo-γ-secretase, we expected that 200 ns are required to reach stable ensembles as evaluated from the C_α_ root mean square deviation (RMSD).(49) The remaining 300 ns provide efficient sampling of the functionally important PS1 helix tilts as expected for this type of motion;(65) these tilts and the associated variations in the catalytic Asp-Asp distance represent the central functional γ-secretase dynamics.

In addition, to understand the conformational dynamics of bound Notch in direct relation to free Notch, the dynamics of the Notch TMD alone in the POPC lipid bilayer was also explored using the same simulation setup for comparison. Three independent simulations of the Notch TMD alone were also run for 500 ns. All simulations were carried out using the high performance computing facility at the Technical University of Denmark during 2018.

### Molecular dynamics data analysis

The trajectory analysis was performed using the GROMACS built-in tools together with FATSLiM for the membrane properties.(66) The membrane thickness, membrane area and membrane area per lipid during were calculated using FATSLiM. The C_α_ RMSD and C_α_ RMSF were calculated with respect to the initial conformation. The hydrogen bonds between the protein side chains and either water or lipid were calculated based on the cutoffs for the angle hydrogen-donor-acceptor and the distance donor-acceptor using default GROMACS set up. The distance between residues was calculated based on the C_α_ atoms. The Gromos method for cluster analysis was used with a RMSD threshold of 0.2 nm.(67) For principle component analysis (PCA), the C_α_ atoms were used for covariance analysis. The obtained eigenvectors represent the direction of collective motion of the atoms and the eigenvalues represent the variation associated with eigenvectors. Typically, more than 90% of the variance is described by less than ten eigenvectors.(68) Visualization was performed using PyMOL, version 2.0.

### Binding energy calculations

Calculations of the binding energy of Notch to γ-secretase were performed using the Molecular mechanics Poisson-Boltzmann surface area (MM-PBSA) method.(69) These energies were computed using the g_mmpbsa package with 100 frames as input, extracted equidistantly from the last 300 ns of each trajectory.(70) For the implicit solvation treatment, dielectric constants of 2 and 80 were used for water and protein, respectively. The binding energy contributions of each residue is the average of the 100 calculations. The obtained binding energy cannot be directly compared to experimental values, and is much too large because it misses compensatory substrate stabilization and entropy. However, the relative contribution of each residue in the three ensembles to Notch binding has these systematic errors canceling to a large extent and is thus more insightful.

**Figure 1.**
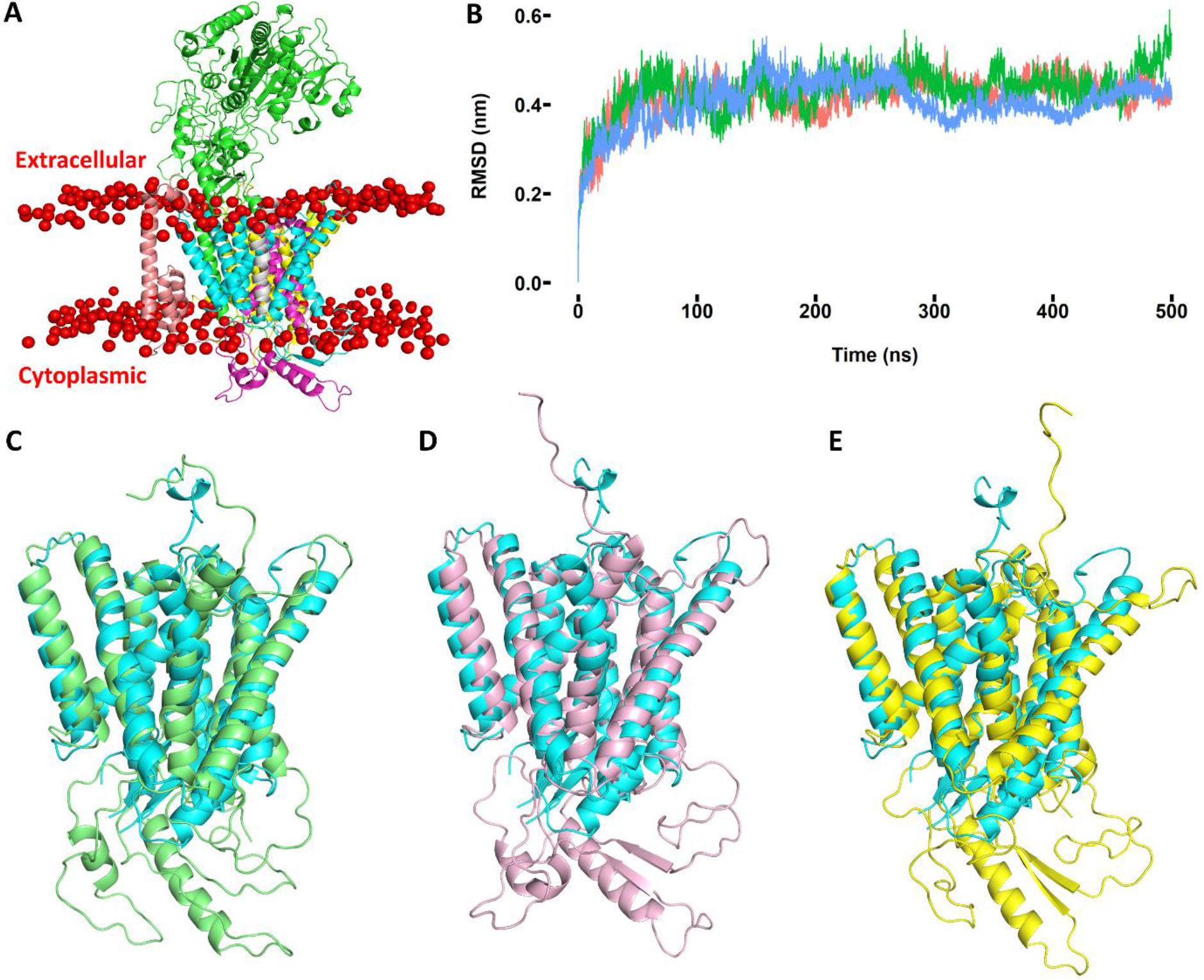
**(A)** Simulated γ-secretase-Notch complex in a POPC bilayer. The red balls represent the head groups of the POPC. The green, cyan, purple, yellow, pink and gray subunits represent NCT, PS1-NTF, PS1-CTF, APH-1, PEN-2 and Notch TMD. **(B)** The C_α_ RMSD (in nm) of three independent γ-secretase-Notch simulations relative to the initial structure used. The red, green and blue lines represent the RMSD for replicate 1, replicate 2 and replicate 3. **(C)** Structural superimposition of cryo-EM structure (PDB: 6IDF, cyan) and the representative structure (green) from simulation 1. **(D)** Same as (C) for simulation 2 (pink). **(E)** Same as (C) for simulation 3 (yellow) (RMSD = 2.07 Å, 2.39 Å and 2.66 Å).

## Results and discussion

### γ-secretase-Notch complex in a POPC bilayer

Among the many identified substrates for γ-secretase, APP-C99 is most extensively studied as cleavage of this substrate directly leads to the Aβ peptides that are considered central to AD.(71) The new cryo-EM structure of Notch-100 bound to γ-secretase(46) provides a test and comparison for our computational approach, which was made before this structure became available using only insights from the separate apo-γ-secretase and Notch TMD structures. The MD simulations provide the all-atom dynamics inside a membrane at ambient temperature. Specific temperature effects can cause protein expansion and increase atomic motions, as quantified by Debye-Waller factors.(72, 73) Since such motions are central to γ-secretase function, the MD simulations strongly complement the cryo-EM data.

The resulting model from our docking of Notch to γ-secretase is shown in Figure 1A. **Figure S1** shows the zoom-in on the PS1 part of the protein complex. The binding pocket involves three transmembrane helices of PS1 (TM2, TM3 and TM5), in agreement with the cryo-EM structure of the γ-secretase-Notch complex.(46) Previous substrate-binding experiments suggested that the TM2, TM6 and TM9 in PS1 were involved in C99 binding, forming an extended surface.(74–76) TM9 may aid initial substrate binding and perhaps represents part of the potential secondary binding site outside the active site of γ-secretase.(77)

The overall dynamic evolution of the three independent trajectories is summarized in Figure 1B, which show the overall C_α_-RMSD for the whole protein and for PS1 alone with respect to the initial structure, respectively. The simulations reach stable ensembles after 100-200 ns in accordance with previous simulations of all-atom apo-γ-secretase in the membrane.(49) Therefore, statistics can be meaningfully collected after 200 ns, and the last 300 ns of the trajectory were used for all the MD analysis described in the following.

We aligned our representative structure obtained from cluster analysis with the cryo-EM structure (PDB: 6IDF) for the PS1-Notch complex, as shown in Figure 1C-1E. Our simulated structures are in good agreement the cryo-EM structure and match all the experimental details especially for the TM helices of PS1, with some differences found in the intracellular part of TM6 and TM7, which are disordered and not complete in the cryo-EM structure, and the C-terminal of Notch. The computed total RMSD for our three representative structure are 2.07 Å, 2.39 Å and 2.66 Å, respectively. Since the average resolution of the experimental structure is 2.6 Å, our simulated models capture the topology of the γ-secretase correctly, but additionally provide the membrane context and the all-atom dynamics at ambient temperature, as described below.

**Figure 2.**
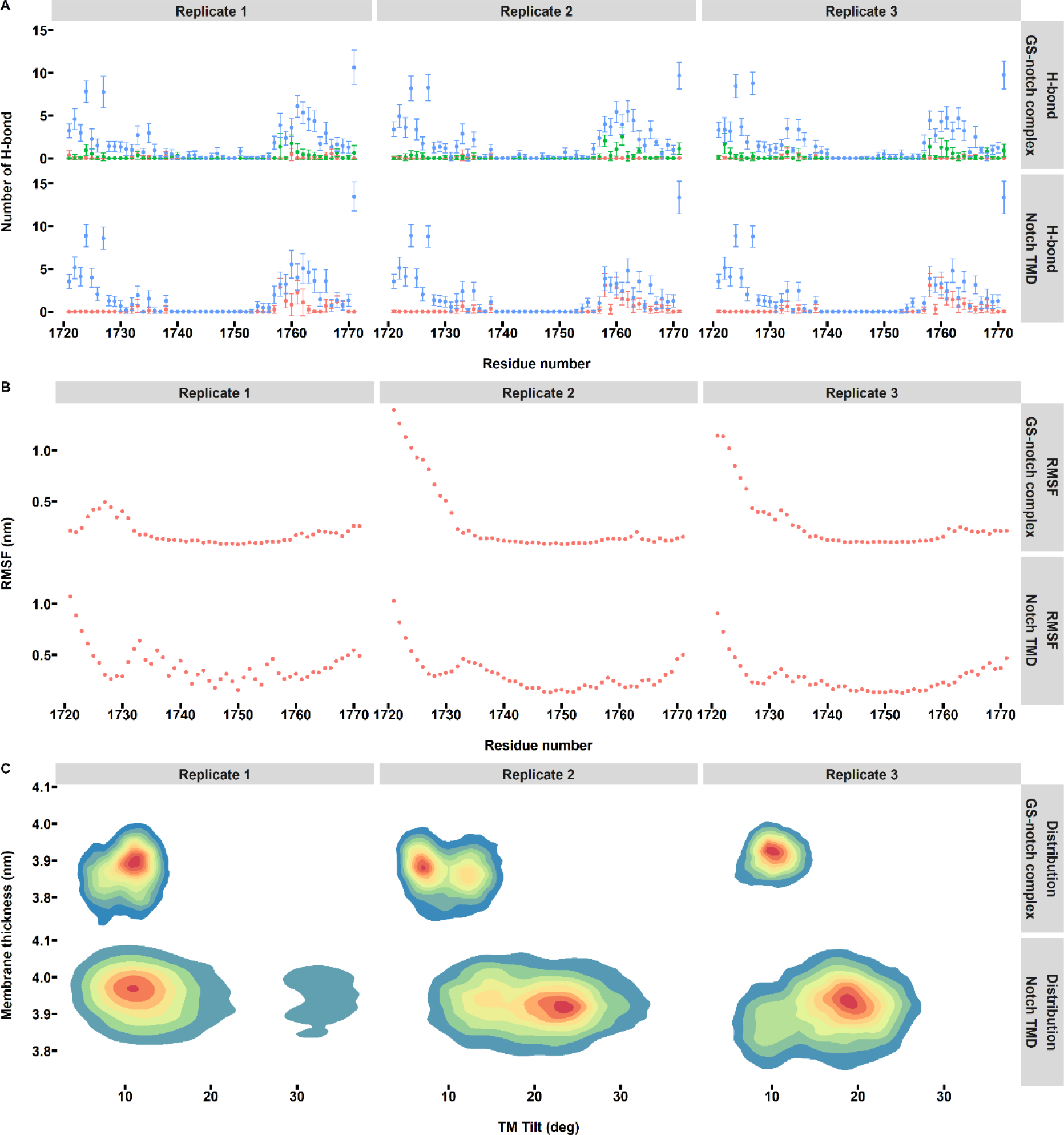
**(A)** Hydrogen bond analysis for free and bound Notch transmembrane domain, showing hydrogen bonds formed with lipids (red), water (blue), and γ-secretase (green). **(B)** The C_α_ RMSF analysis for free and bound Notch transmembrane domain. **(C)** Simulated distributions of free and bound Notch transmembrane domain models projected onto the transmembrane tilt angle and membrane thickness.

In order to ensure the realism of our simulated membrane-protein system, we calculated the membrane order parameters (**Figure S2**), density of the POPC bilayer (**Figure S2**) and several other membrane properties (**Figure S3**). As shown in **Figure S2** and **S3**, the order parameters (S_cd_), which evaluate the flexibility of the lipids as a function of position within the lipid chain, and the membrane properties are in excellent agreement with the typical experimental and MD values for realistic membrane systems.(78, 79) From this analysis, we conclude that our protein-water-membrane system at physiological temperature and ion strength is a realistic model of mature γ-secretase-Notch.

### Dynamics of the Notch transmembrane domain

In order to understand how Notch changes structure upon binding to γ-secretase, we performed all-atom MD simulations of the Notch TMD in the POPC bilayer both in the free form and bound to γ-secretase. We computed the number of the hydrogen bonds formed between either water, lipid or γ-secretase as shown in Figure 2A. In agreement with NMR data,(48, 80) the N-terminal of the Notch TMD was water-exposed and consequently maintained a large number of hydrogen bonds with water during the entire simulations. The residues A1732 and Q1733 located in the membrane interface formed hydrogen bonds with both water and lipid head groups; in the cryo-EM structure,(46) the lipids are not there and this these residues interact with water instead. As shown in Figure 2A, after being totally devoid of water within the transmembrane segment, hydrogen bonding to water was observed again at G1753 and V1754 of the S3 site, which is the initial γ-secretase cleavage site and is in close contact with the two catalytic aspartates in our structures (in the cryo-EM structure, one of these aspartates is mutated to alanine).(48, 80, 81) We propose that we have thus identified the catalytic water required for nucleophilic attack at the S3 site, important because it could be speculated how water reaches the hydrophobic TMD to enable the catalytic process. Local intermediate time-scale motions were also found for V1754 during NMR experiments.(80) The hydrogen bonding network clearly suggests that the cytosolic C-terminal side of Notch dips back into the membrane as the consequence of the LWF (1767-1769) motif. This is fully in line with previous NMR data suggesting that the membrane reentrant segment centers at the LWF motif.(48) Such dynamics cannot be seen in the cryo-EM structure because it does not feature a membrane.

Compared to free Notch TMD, the bound state had its C-terminal side mainly interacting with γ-secretase instead of membrane lipids. In addition, the hydrogen bonds formed at the beginning of Notch TMD (QSETVE 1722-1727) indicate interactions between these residues and γ-secretase, as is also visible in the new cryo-EM structure of the Notch-γ-secretase complex.(46) The atomic motions of Notch as measured by C_α_ RMSF are shown in Figure 2B. High fluctuations were mainly found at the N-terminal side, in agreement with NMR experiments suggesting that the N-terminal side is particularly flexible.(80) We identify less fluctuation at the C-terminal side showing that this side is more ordered.

The membrane thickness affects the topology of the overall protein-substrate complex and is of importance for regulating Notch cleavage by γ-secretase.(48, 82) According to the FIST (Fit-Stay-Trim) model,(43, 83) the more compact (semi-open) state of γ-secretase favors stronger substrate binding and retention time and maximal trimming to shorter products, whereas PS1 mutations or thinner membranes will favor the open state, which has a shorter residence time of substrates and thus produces less cleavage products, with relatively more of the long products. To understand this crosstalk between membrane and protein dynamics, we computed the projection of the MD simulations onto the membrane thickness and Notch TMD tilt angle, as shown in Figure 2C. The free state of Notch TMD displayed much more tilting relative to the membrane which has been observed before,(48) whereas the bound Notch state was much more rigid due to the interactions with γ-secretase.

The variation in the membrane thickness was particularly interesting and shows clearly that the membrane-protein system is highly dynamic and work together to produce the conformational changes in the protein and substrate that define the open and compact states, as we discuss further below. These findings strongly support the hypothesis that the membrane dynamics play a central role in γ-secretase cleavage(84, 85) and makes the complementary study of MD membrane-protein dynamics very important as a supplement to the cryo-EM data. The most notable differential effect of free and bound Notch is that the Notch TMD alone retained its α-helicity during the full simulations, whereas in the bound state the helix has a tendency to lose helicity for some specific residues, notably 1742-1743 and 1752 at the S3 site (**Figure S5)**; thus the membrane dynamics support a helix destabilizing effect upon substrate binding which was indicated from the cryo-EM data and which may aid in exposing the peptide bonds to hydrolytic cleavage by the aspartates.(46, 86)

### Distinct catalytic site dynamics affect space and Notch positioning

The dynamic variation in the distance between the catalytic aspartates and the Notch cleavage sites are of special relevance to Notch cleavage. These distances and their dynamic variations are shown in Figure 3A. These variations are not visible in the cryo-EM data but probably cause the imprecise cleavage leading to several NICDs.(47)

In our simulations the catalytic D385 was closer to the cleavage site at V1754 (Figure 3A, blue). This is in line with a previous hypothesis that the protonated D385 can donate an initial proton to the substrate, with a short distance enabling the proton transfer.(87) The D257-D385 distance defines the space available in the catalytic site and was thus emphasized in previous cryo-EM studies; in the apo-structure from 2015, the distance was 10.6 Å.(88) In the new Notch-bound structure,(46) the corresponding mutant Ala-Asp distance is 10.4 Å. Interestingly, this distance was markedly shorter in replicate 3, indicating a more compact substrate-bound ensemble, and this shorter distance also produces a shorter distance between the catalytic D257 and the cleavage sites. The presence of compact and looser substrate bound states do not emerge from the static average cryo-EM electron density maps, yet these different states are essential for explaining the different outcomes of substrate cleavage, as analyzed further below.

**Figure 3.**
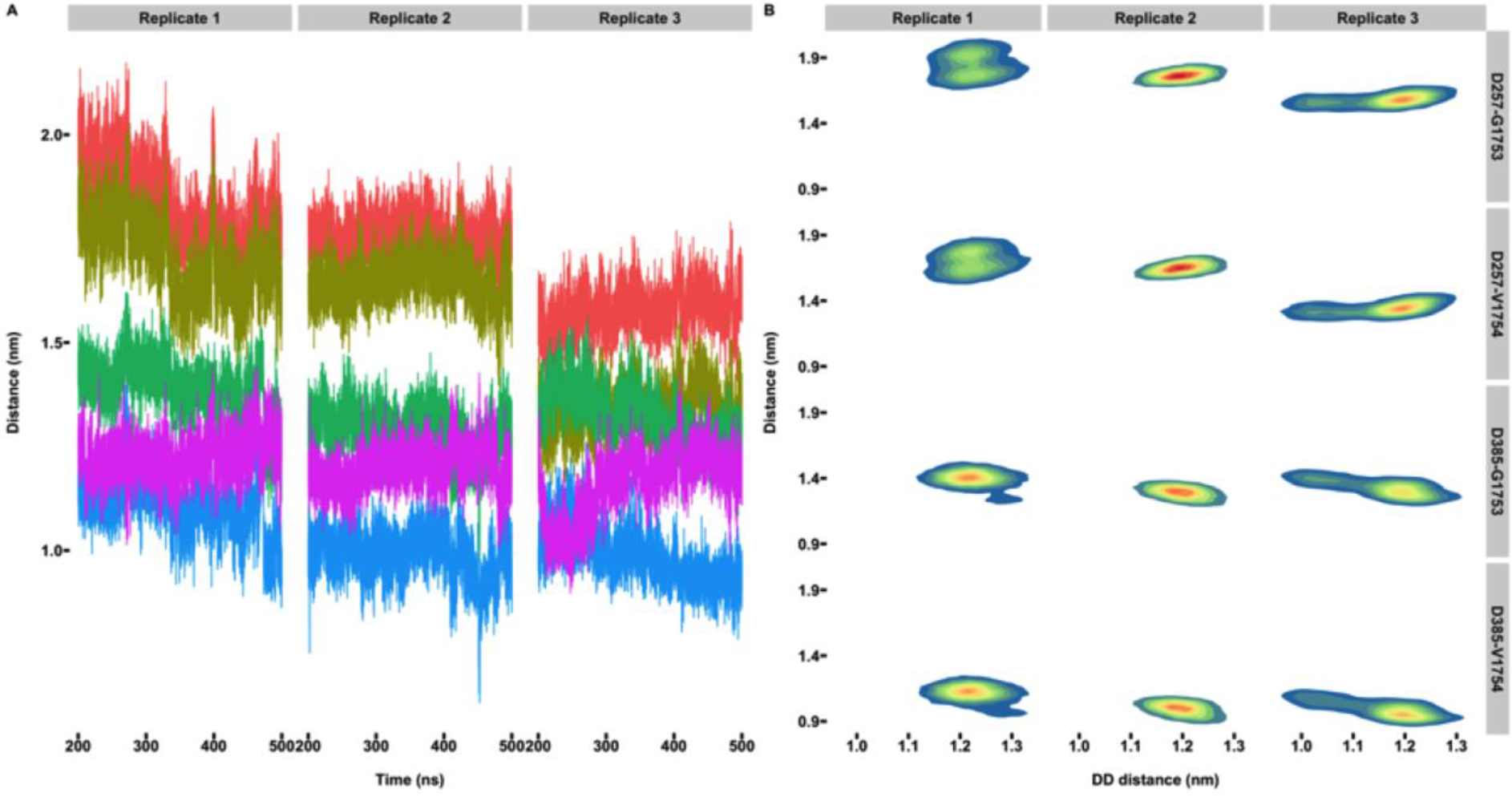
**(A)** Simulated distance between PS1-D257 and Notch-G1753 (red), PS1-D257 and Notch-V1754 (yellow), PS1-D385 and Notch-G1753 (green), PS1-D385 and Notch-V1754 (blue), and PS1-D257 and D385 (purple). **(B)** Distributions of the distance between PS1-D257 and D385 (DD) and their distance to residues of the Notch cleavage site.

As seen from Figure 3B, proximity to one cleavage site associates with increased distance to the other cleavage site in a ping-pong fashion. The dynamics of this relationship suggests that the bound substrate orients itself after initial cleavage by aligning the next peptide bond towards the opposite catalytic aspartate in a ping-pong cleavage mechanism. The peptide bonds with a separation of three amino acids will be approximately exposed to each of the two aspartates, because of the helix winding, consistent with the tree-amino acid spaced cleavage patterns that are most commonly observed.(15)

Membrane thickness may affect substrate cleavage, with thicker membranes favoring more extensive substrate trimming, leading to more short products,(89, 90) consistent with the FIST mechanism.(43, 49, 83) An increase of carbon numbers in the lipid chain accordingly also increases γ-secretase activity.(84) Inspired by this finding, we computed the correlation between membrane thickness and the important catalytic distances (**Figure S4**). As seen from **Figure S4**, we found a relatively thicker membrane in replicate 3 which also displayed a shorter D257-D385 distance, confirming the presence of a more compact state in this ensemble. Previous MD simulations of γ-secretase indicated conformational changes of PS1 in response to hydrophobic mismatches in the lipid bilayer.(89)

Our computed tilt angle distributions of TM helices in PS1 (**Figure S6)** show marked differences between the simulations, with replicate 2 giving a distinctly variable two-state behavior for several TM helices representing the more open state, whereas replicate 1 and 3 are less variable. Not only the variation in tilt angles but also the prevalent tilt angle of each TM helix varies between the ensembles; we expect the cryo-EM data to reflect an average of these motions. These differences largely define the open and compact states central to the FIST mechanism as seen in previous MD simulations also of apo-γ-secretase in the membrane.(49) Compared to non-substrate bound γ-secretase, the substrate-bound protein was more rigid as indicated by all tilt movements shown in **Figure S6**. The TM2, TM6 and TM9 are involved in initial substrate binding.(77) The marked change in Notch structure upon binding (**Figure S6**) supports the notion that the membrane substrates of γ-secretase undergo conformational rearrangement during the cleavage process.(46)

The overall atomic motility as quantified by the RMSF in **Figure S7** confirmed the high mobility of TM6 and TM7 of PS1. Among the other subunits, NCT displayed the typical large fluctuations consistent with its gatekeeper role in substrate recognition;(91) these breathing modes of NCT have been seen routinely in MD simulations.(49),(87) Moreover, as shown in **Figure S7**, the APH-1 and PEN-2 subunits exhibited relatively more rigid dynamics with lower RMSF values, and the residues in PS1 close to these subunits also showed low mobility (**Figure S7**).

We correlated the simulated PS1 TM helix tilt angles and different distance profiles as shown in **Figure S8**. Most distributions indicate correlation with Notch TMD tilt modifications suggesting the importance of this particular movement of Notch vs. other possible movements of the substrate. In addition, other notable changes were observed: (1) The distance between D257 and the Notch cleavage site showed two distinct states; the shorter distance correlated with shorter D257-D385 distance; (2) the distance between D385 and the Notch cleavage site correlated with the tilting of the Notch TMD and the tilting of TM6 of PS1; in replicate 1, TM6 and Notch tilting both correlated with the D257-G1753 and D257-V1754 distances; (3) for the D257-D385 distance alone, no correlation with specific helix tilt movements were found in contrast to previous non-substrate bound MD simulations.(49)

**Figure 4.**
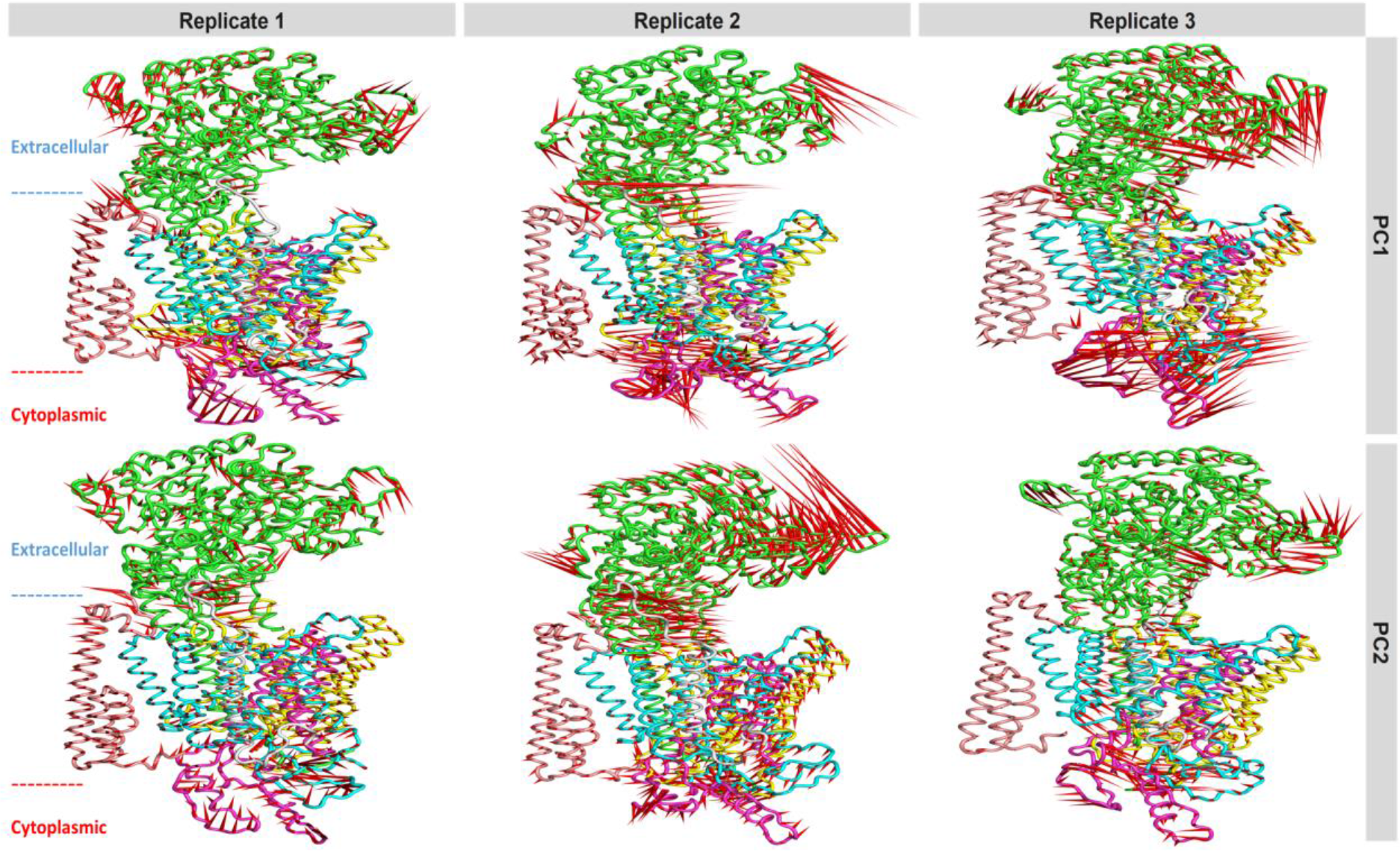
Porcupine plots representing the collective motions of Notch-γ-secretase within the membrane along the first and second principal components.

**Figure 5.**
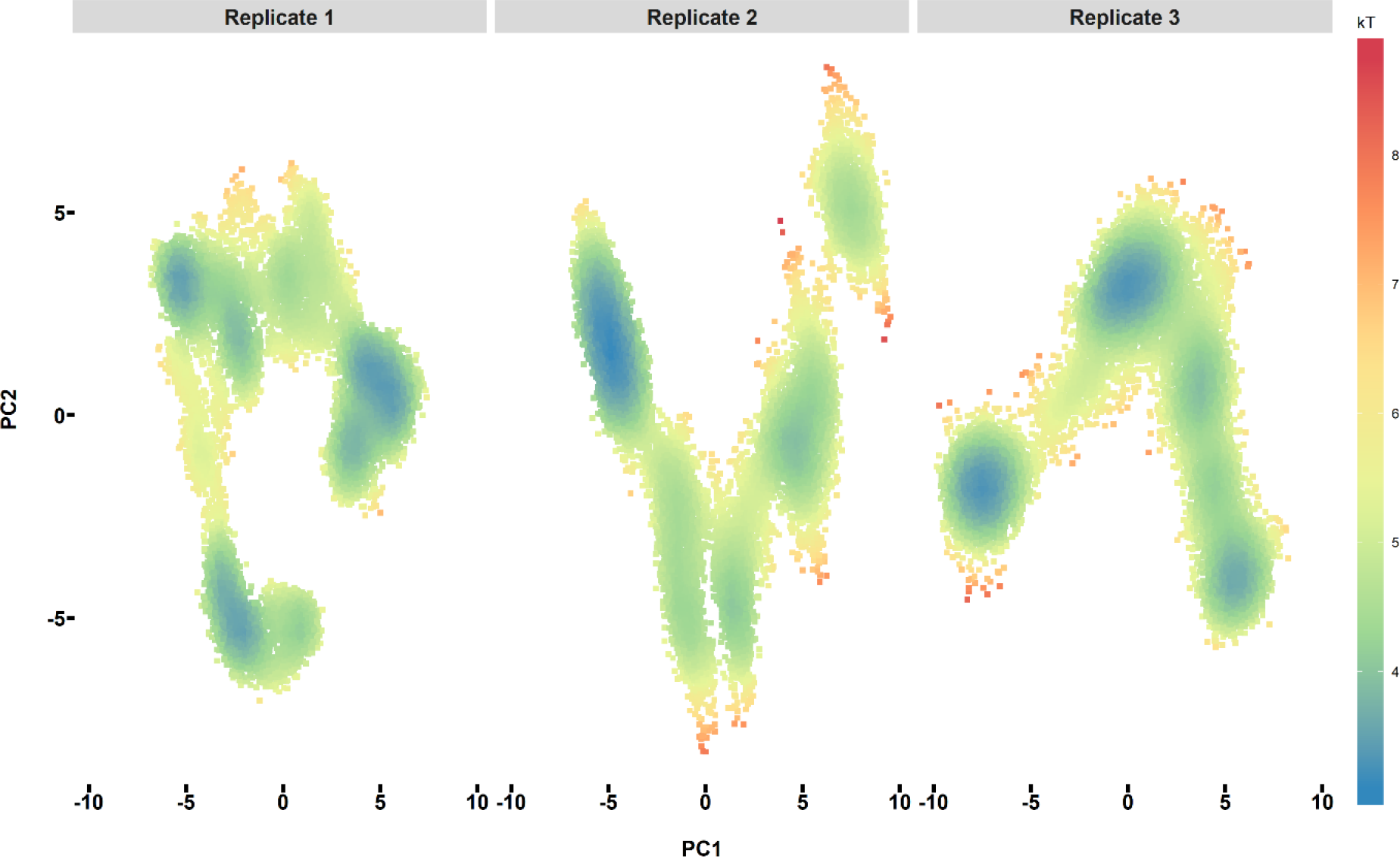
Free energy landscape of the Notch-γ-secretase complex projected onto the first and second principal components PC1 and PC2 obtained from principal component analysis. The free energy is plotted to the right with lower energy colored blue.

### Global conformation states of the Notch-γ-secretase complex in the membrane

To understand the motions of the full protein-substrate complex with the membrane in more detail, porcupine plots along the direction of PC 1 and PC 2 were plotted as shown in Figure 4; they accounted for the majority (24.4%, 31.6% and 30.6% for replicate 1, replicate 2 and replicate 3, respectively) of all motions. As seen from Figure 4, there are three very predominant motions: (1) The extracellular NCT showed up/down movement relative to the membrane indicating its gatekeeper role in substrate recognition in agreement with previous experimental data;(92) (2) the N-terminal of the Notch TMD exhibited movement towards and away from the hydrophilic loop 1 of PS1 which is in line with previous suggestions that this loop is also involved in substrate recognition;(93, 94) (3) interestingly, another clear motion was identified in the cytosolic part of TM6 and TM7 of PS1. In the recent published cryo-EM structure of the Notch-γ-secretase complex, this part of PS1 changes structural rearrangement relative to the apo structure, as quantified by our collective atomic motions.(46) Taken together, the results suggest that NCT and the hydrophilic loop 1 interact with the N-terminal of Notch TMD, and the movement of cytosolic parts ofTM6 and TM7 of PS1 introduce conformational changes to the substrate-bound state.

To summarize the ensembles of the membrane protein, we performed free energy landscape analysis based on the PCA at 303 K as summarized in Figure 5. The free energy is defined by kT units according to the conversion ΔG = −kTln*P*(PC1, PC2), where *P* is the probability (prevalence during simulation) of a given coordinate of PC1 and PC2. The landscapes did not feature any deep free energy wells but rather display multiple shallow minima indicating that the Notch-γ-secretase system in the membrane is approximately a three-state system. This observation rationalizes the disorder seen in cryo-EM data(49) and provides specific explicit atom components to these motions.

To understand the dominating conformation states in structural detail, we performed cluster analysis to extract the top-3 most visited structures using a threshold of 0.2 nm. The alignment of the extracted structures from each replicate simulation is shown in Figure 6. A recent cryo-EM study of the Notch-γ-secretase(46) complex brings new insights into Notch cleavage by γ-secretase and has implications also for our study. According to this study, Notch binding is facilitated by the formation of a hybrid β-sheet between the Notch TMD (β3) and two substrate-induced strands in PS1 (β1 and β2), leading to remarkable structural changes in both structures that orient the cleavage site of Notch towards the PS1 catalytic residues.(46) A partial unzipping of the Notch TMD was argued to be visible from the cryo-EM electron density maps.

**Figure 6.**
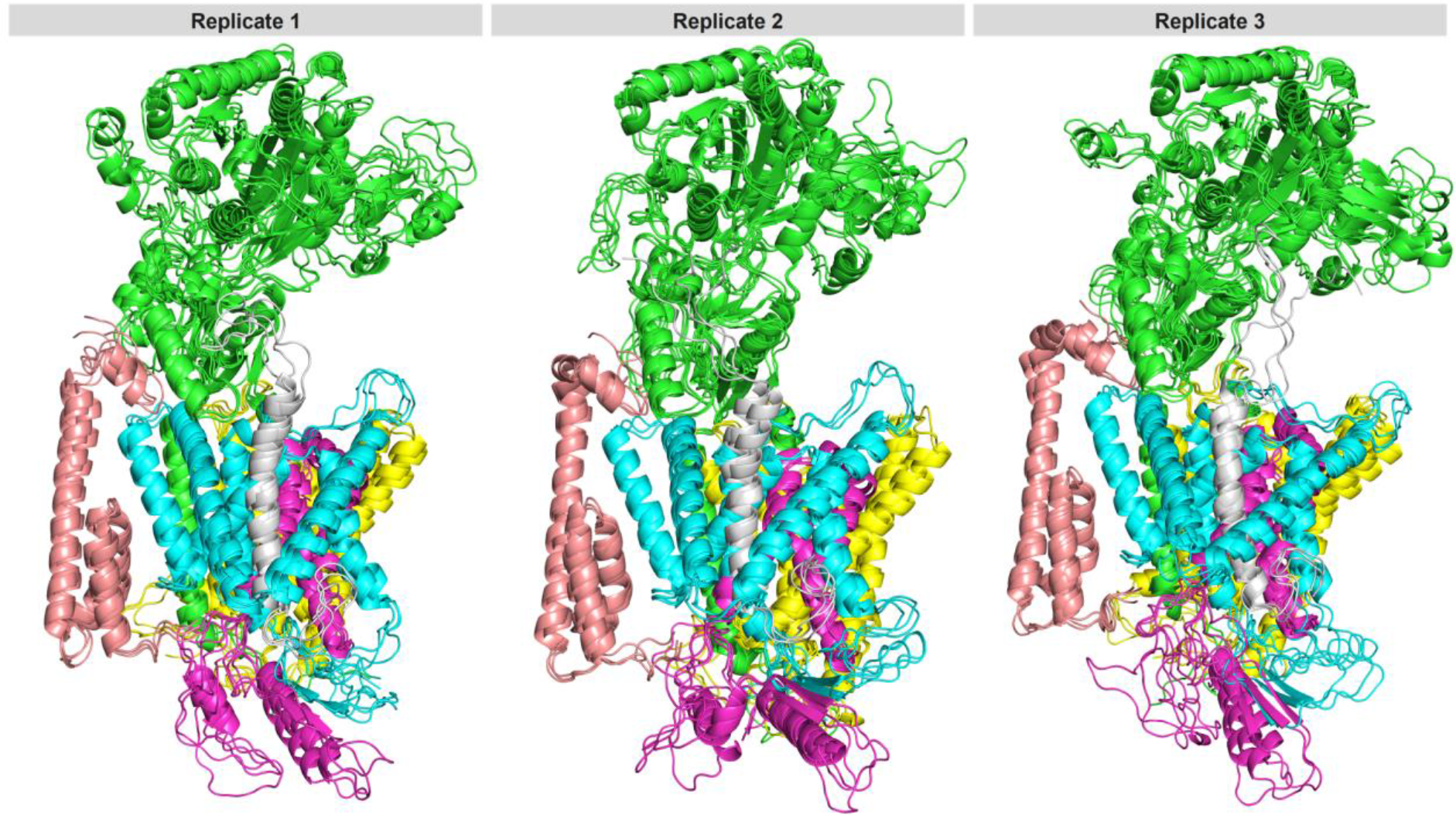
Alignment of the top-3 representative structures obtained from cluster analysis. The green, cyan, purple, yellow, pink and gray subunits represent NCT, PS1-NTF, PS1-CTF, APH-1, PEN-2 and the Notch TMD, respectively.

It is of substantial interest to know if these features, while very appealing as mechanistic explanations for Notch cleavage, are affected by the membrane at ambient temperature where the conformations of the transition-prone HL1 and HL2 may change. Deletion of residues involved in forming the β1 or β2 in PS1 was found to abolish proteolytic activity, which indicates that the residues, and perhaps also the β-sheet, are important for γ-secretase activity.(46) The stable Notch-γ-secretase complex was obtained via covalent crosslinking by introducing a cysteine residue in PS1 and mutation of the catalytic D385 to alanine to prevent turnover during sample preparation; thus the cryo-EM structure represents an inactive state; however, the reason we see several substrate-bound states is more likely due to the fact that the cryo-EM shows an average snapshot of the rapidly frozen sample, and thus does not reveal the conformational changes specifically. Our identified conformation states in the membrane with distinct distances to the cleavage sites (Figure 3) may explain the imprecision of S3 cleavage and the different cleavage patterns,(41) which cannot be explained from one static average structure.

As can be seen from Figure 6, we did observe two β-strands in PS1 in all the representative structures except the top-1 structure of replicate 1. For β1 formed in TM6 of PS1, we observed the exact same residues (288-290) forming this β-sheet in all relevant structures. The first discovery of strand motifs in PS1 was from an MD simulation of PS1 alone,(95) which has now also been seen experimentally,(46) but our simulations show that this tendency is quite persistent also in the full complex. A notable difference between our simulated structures and the cryo-EM structure is a residue shift for β2 formed in TM7 of PS1 which were 328-331 in our simulations instead of 377-381 determined for the cryo-EM structure. In addition, as illustrated in Figure 6, we did not observe the hybrid β-sheet via a strand in Notch.

It was suggested that the transmembrane substrate helix of intramembrane proteases unwinds during binding to enable the catalytic aspartates to access the peptide bonds,(96) and solution NMR chemical shifts indicate that this is also the case for cleavage of C99.(97) Such a mechanism requires helix-destabilizing residues to partially unzip during substrate binding. The recent cryo-EM structure is claimed to show such a partial unwinding within the Notch-TMD.(46) Motions of this type are expected in the dynamics of any helix, and thus it is relevant to know if this tendency is distinct for bound Notch in comparison to Notch free in the membrane as a control. To determine this, we calculated the number of Notch residues associated with α-helix during the last 300 ns; the results are shown in Figure 7. As a good reference point, the α-helix tendency of the TM helices of PS1 was quite stable. In marked contrast, there was a considerable difference in helix tendency for free and bound Notch TMD in replicate 2 and 3.

The loss of helicity is not accompanied by β-sheet formation in our simulations possibly because we use a smaller Notch model, or because the mutated Notch binds differently in some minor aspects, or because of the membrane. Cryo-EM experiments used Notch-100 (1721-1820), which has a notable intracellular part that could interact with cytosolic parts of TM6 and TM7 of PS1 and then undergo the coil-β-transition. Due to lack of templates and to avoid bias, we used Notch-51 (1721-1771); the C-terminal juxtamembrane part dipped back to the membrane leading to lack of interaction between Notch and the cytosolic TM6 and TM7 of PS1 thus preventing β-sheet formation. β-sheets formed in TM6 and TM7 of PS1 were first observed in our previous MD simulations of apo-γ-secretase(49, 95) providing support for the role of β-strands in γ-secretase PS1 dynamics; it is instructive to see that cryo-EM data can confirm these findings, which exemplify the realism of our general protocols.

### Mechanism of imprecise substrate cleavage by two-state γ-secretase

Cryo-EM data of substrate bound γ-secretase reveal one main state; yet two states are required to explain the cleavage patterns according to the FIST model.(43, 49, 83) According to this model, the protein-substrate system has two conformation states, a looser (open) and a more compact (semi-open), defined by the TM2, TM3, and TM6 tilts of PS1. These tilts reflect squeezing of the substrate. The more compact state has higher substrate affinity, keeps the substrate predominantly at the site that leads to more of the short products, it keeps the substrate for longer and “squeezes” it to shorter products at higher overall activity. In contrast, the open (loose) state with longer distance between the aspartates and cleavage sites gives a mixture of cleavage initiation sites due to imprecise positioning, and displays weaker binding, leading to earlier loss of products at longer length.(43, 49)

Our simulations reveal the presence in the membrane ensemble of a compact and looser state of Notch binding which differ notably in the catalytically important distances between the aspartates and S3 sites. The distances are correlated such that closeness to one site increases the distance to the other, and the dynamics indicate residues at the S3 site at the membrane interface ping-pong movements that adjust the cleavage sites to the aspartates. The catalytic water that we also claim to have identified is in direct contact with the S3 site in these dynamics, whereas no water is found inside the membrane elsewhere. Our ensembles indicate that imprecise S3 cleavage can occur for Notch as for C99 via compact and loose states, which would explain the existence of multiple NICDs formed.(40)

We calculated the energy contribution of each residue to Notch binding (calculated using MM-PBSA) as shown in **Figure S9-S11**. The contributing residues were generally identical in all the replicates. The C-terminal of PS1 contributed substantially to the Notch binding, which is in line with a previous suggestion that the CTF of PS1 is most important for substrate binding. In addition, we note that the positively charged motif (RKRRR 1758-1762) of Notch contributed significantly to overall binding and thus serves as a “membrane anchor” of Notch that prevents its displacement towards the extracellular side. Previously, the positively charged lysine-53 in C99 was found to be critical for C99 binding.(24)

We also noticed that the residues in PS1-CTF that contribute considerably to binding are all charged amino acids (mainly negatively charged Asp and Glu). In our previous saturation mutagenesis study on PS1 related to FAD, we identified many mutations of lipid-exposed charged residues that cause FAD.(83) The present study suggests this may due to their involvement in substrate binding, and the mutations could reduce the electrostatic binding affinity leading to more loose substrate binding, giving more of both trimming pathways and earlier substrate release of longer products, notably giving increased Aβ_42_/Aβ_40_ ratios that correlate with disease severity of the PS1 mutations.(25, 26)

**Figure 7.**
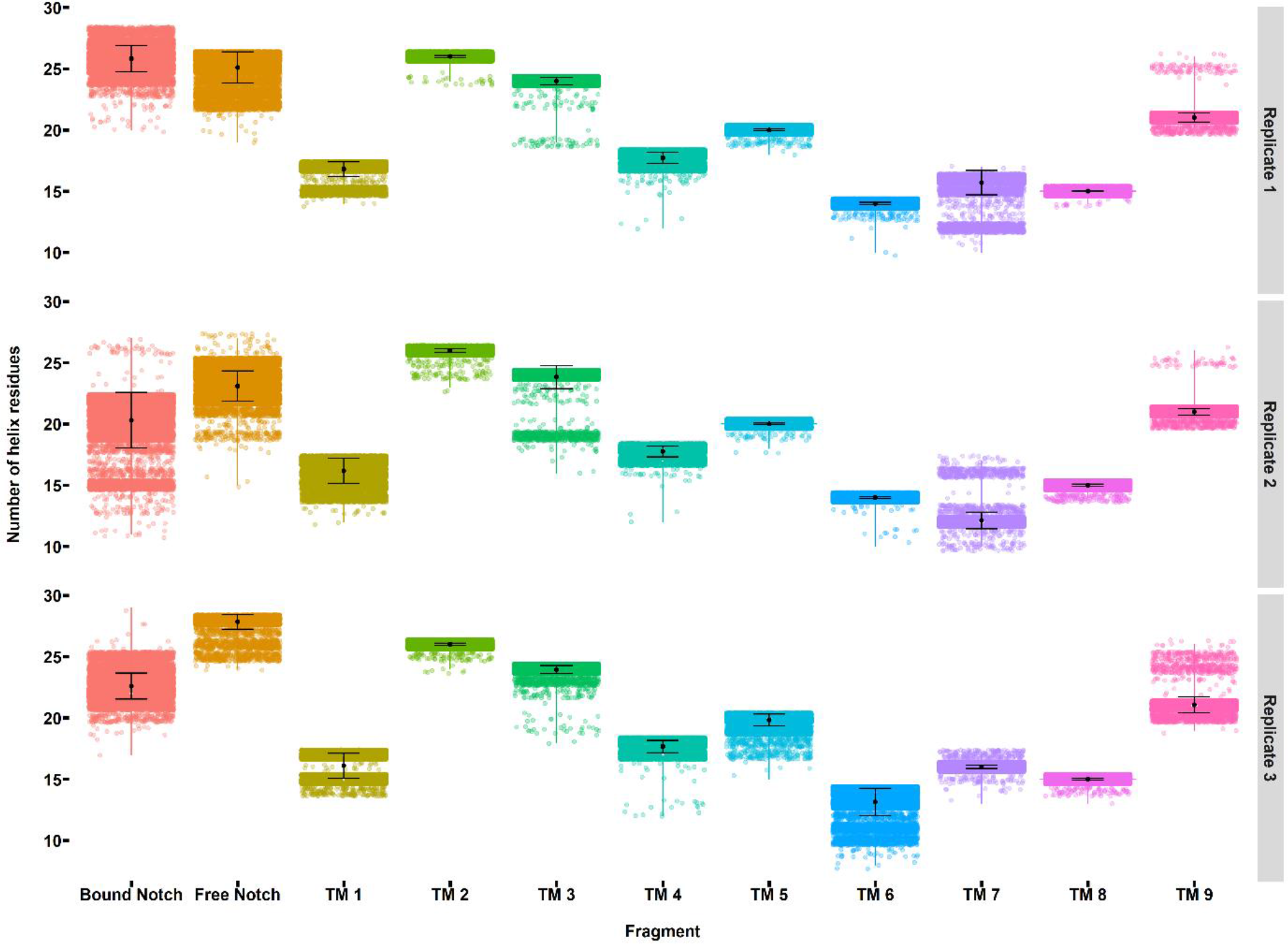
The distribution of residues associated with α-helix in free and bound Notch TMD and in the nine transmembrane helices of PS1 during the last 300 ns simulation of our three independent simulations.

## Conclusions

Our MD simulations have elucidated the detailed atomic dynamics of the Notch-γ-secretase complex in a membrane, which substantially complement the cryo-EM structure of Notch bound to γ-secretase.(46) Our structural ensembles developed from apo-enzyme models are in essential agreement with the cryo-EM data (RMSD ~2.07-2.66 Å vs. average experimental resolution of 2.6 Å) and more importantly provide the all-atom motions of the full protein-membrane system at normal temperature (303 K).

We show by direct comparison to similar simulations of free Notch also in a membrane that Notch loses critical α-helicity in the middle of the Notch TMD due to low helix propensity of 2-3 residues. mostly to the N-terminal side of the S3 cleavage site. These residues are thus probably responsible for inducing a partial substrate unwinding as suggested previously(96) and supported by NMR data,(97) and as recently indicated from the cryo-EM data.(46) We also show that the membrane thickness plays an important role in modulating the catalytically important residue distances, thereby providing a mechanistic basis for the effect of membrane composition on Notch cleavage.

We consistently observe β-strand formation in several exposed parts of PS1, in agreement with previous studies in our group.(49, 95) Cryo-EM data have also recently supported the presence of β-strand formation in PS1.(46) However, our simulations do not indicate a specific hybrid β-sheet being formed in the membrane, which can be an artifact of the simulations, or possibly due to the absence of the membrane in the cryo-EM data. We also find that the RKRRR motif (1758-1762) contributes significantly to Notch binding (MM-PBSA calculations) and serves as a “membrane anchor” that prevents Notch displacement toward the extracellular side of the membrane. We also observe that while water is totally excluded from the TMD region, water can transiently hydrogen bond to G1753 and V1754 in our structures; we suggest that this water is the catalytic water acting as the nucleophile during peptide bond cleavage.

Our most important finding is that Notch binding induces three conformation states, including two that differ in the catalytic site size: One state is more open and the other is more compact. Such states are not seen in the average cryo-EM density but are required for explaining the different outcome of cleaving Notch by γ-secretase. Specifically, the sloppy cleavage of the more open state enables multiple cleavage sites around S3 to be attacked in a ping-pong fashion, leading to more different products as seen from assays.(40, 41) These findings are in agreement with the FIST model(43, 49, 83) where the loose “grab” causes imprecise cleavage at several sites and releases substrates earlier with longer average length. Our simulations suggest that Notch undergoes qualitatively similar “sloppy” proteolysis as C99 due to similar involvement of loose and compact substrate-binding states. We suspect that any selective targeting of C99 over Notch will require an emphasis on these conformation states in the membrane rather than the average densities seen at cryo temperature, and thus our structural dynamics should be of value to the development of selective drugs targeting C99 cleavage over Notch cleavage.

## List of abbreviations

Aβ: β-amyloid
AD: Alzheimer's disease
APH: Anterior pharynx defective
APP: β-amyloid precursor protein
C83/C99: C-terminal fragment of β-amyloid precursor protein
Cryo-EM: Cryo-electron microscopy
CTF: C-terminal fragment of PS1
FAD: Familial Alzheimer's disease
FATSLiM: Fast analysis toolbox for simulations of lipid membranes
FIST: Fit stay trim
GPU: Graphics processing unit
GROMACS: Groningen machine for chemical simulations
HL: Hydrophilic loop
LINCS: Linear constraint solver
MD: Molecular dynamics
MMPBSA: Molecular mechanics Poisson-Boltzmann surface area
NCT: Nicastrin
NICD: Notch intracellular domain
NMR: Nuclear magnetic resonance
NPT: Constant temperature, constant pressure ensemble
NTF: N-terminal fragment of PS1
NVT: Constant temperature, constant volume ensemble
PC: Principle component
PCA: Principle component analysis
PEN-2: Presenilin enhancer 2
PME: Particle mesh Ewald
POPC: 1-palmitoyl-2-oleoyl-glycero-3-phosphocholine
PPM: Position protein in membrane
PS: Presenilin
RMSD: Root mean square deviation
RMSF: Root mean square fluctuation
TM: Transmembrane helix
TMD: Transmembrane domain

## Authors’ contributions

NT and KPK designed the work. NT and BD performed all computations. NT and KPK analyzed the data. NT, BD, and KPK wrote the paper. All authors read and approved the final manuscript.

## Acknowledgements

The authors acknowledge computer time from DTU high-performance computing facility, Lyngby, DTU. The Danish Council for Independent Research | Natural Sciences (DFF), grant case 7016-00079B, and the Novo Nordisk Foundation, grant NNF17OC0028860, are gratefully acknowledged for supporting this work.

